# A lipid-free and insulin-supplemented medium supports *de novo* fatty acid synthesis gene activation in melanoma cells

**DOI:** 10.1101/479964

**Authors:** Su Wu, Anders M. Näär

## Abstract

While investigating the role played by *de novo* lipid (DNL) biosynthesis in cancer cells, we sought a medium condition that would support cell proliferation without providing any serum lipids. Here we report that a defined serum free cell culture medium condition containing insulin, transferrin and selenium (ITS) supports controlled study of transcriptional regulation of *de novo* fatty acid (DNFA) production and *de novo* cholesterol synthesis (DNCS) in melanoma cell lines. This lipid-free ITS medium is able to support continuous proliferation of several melanoma cell lines that utilize DNL to support their lipid requirements. We show that the ITS medium stimulates gene transcription in support of both DNFA and DNCS, specifically mediated by SREBP1/2 in melanoma cells. We further found that the ITS medium promoted SREBP1 nuclear localization and occupancy on DNFA gene promoters. Our data show clear utility of this serum and lipid-free medium for melanoma cancer cell culture and lipid-related areas of investigation.

## Introduction

*de novo* lipid (DNL) synthesis is the metabolic pathway that converts carbohydrates into fatty acids, cholesterol, phospholipids, triglycerides, and other cellular lipids required for normal cellular homeostasis and proliferation/growth. In healthy adults, DNL is for the most part restricted to liver and adipose tissues for energy storage or distribution to other tissues. Many malignant cancer cells also exhibit elevated DNL as a hallmark adaptation to support proliferation and survival (1, 2). Among the DNL pathways, *de novo* fatty acid (DNFA) biosynthesis is of particular interest as a potential therapeutic target for cancers (3, 4). DNFA is primarily regulated at the mRNA level of catalytic enzymes that drive the biosynthetic reactions (5), a transcriptional process under the control of the sterol regulatory element-binding protein 1 (SREBP1) (6). However, DNFA gene regulation is still not fully understood at the molecular level. Cell culture studies of DNFA with widely-accepted serum-containing medium conditions are often confounded by the presence of external lipids, with consequent difficulty to disentangle the respective effects of lipid synthesis and lipid update, both of which may occur even among cells that are able to survive and proliferate using DNFA alone (6, 7).

Most cell culture media are composed of serum supplements and basal media (BMs), each of which may contain confounding factors for DNFA studies. Serum typically contains external lipids in two forms: non-esterified free fatty acids (FFAs) associated with albumin, and lipoproteins that carry triglycerides, cholesterol and phospholipids encapsulated by apoproteins (8, 9). Cells can take up FFAs *via* physical diffusion across the cellular membrane (10) or *via* active transport aided by membrane-associated protein CD36 or FATPs (11). Lipoprotein uptake occurs through binding to cell surface receptors (e.g. LDL receptor; LDLR) and the bound lipoprotein/receptor complex then undergoes endocytosis, intracellular/lysosomal release of the cargo and receptor recycling to the plasma membrane (12). After transport into cells, both FFAs and lipoprotein lipids are potent DNFA inhibitors in a classic negative feedback manner (13). Intracellular cholesterol interferes with trafficking of SREBP1 from the ER to the Golgi apparatus – an essential step in post-translational processing and maturation of SREBP1 – and consequently inhibits DNFA enzyme expression (6). Polyunsaturated FAs (PUFAs) have also been observed to inhibit both SREBP1 mRNA transcription and protein maturation, resulting in decreased expression of DNFA enzyme genes (14–16). DNFA expression studies have often employed lipoprotein-deficient serum (LPDS) to limit negative feedback by serum lipids. LPDS is free of lipoproteins, which are removed by ultracentrifugation, yet retains FFAs – which are confounding for DNFA studies (17). An alternative is delipidated serum (7), although that is prepared by organic solvent extraction and is not completely lipid-free (18). Preparation protocols vary widely in organic solvent composition and extraction time, and quality variation between batches is common (19, 20).

Basal media (BM) often contain glucose, which besides its bioenergetic role as fuel for ATP synthesis also serves as a carbon source for biosynthesis of amino acids, nucleotides, other carbohydrates, and lipids. Two of the most common BMs for cancer cell culture are Roswell Park Memorial Institute (RPMI) 1640 medium (21), with 11.11 mM glucose concentration (normal glucose level); and Dulbecco’s modified Eagle’s medium (DMEM), which contains 25 mM glucose (high glucose level). In cultured hepatocytes and adipocytes, high glucose conditions stimulate lipogenesis and DNFA gene expression (22, 23). Healthy livers employ glucose for DNFA principally by glycolytic citrate generation and the tricarboxylic acid (TCA) cycle with oxidative phosphorylation (24, 25). Glycolysis produces pyruvate that mitochondria metabolize into citrate and ATP in the TCA cycle. The citrate then translocates to cytosol where it is cleaved by ATP-citrate lyase (ACLY) to produce acetyl-CoA as a substrate for DNFA (26–28). In contrast to healthy livers, tumors display enhanced glycolytic activity but impaired oxidative phosphorylation. Pyruvate is often metabolized to lactate within the cytoplasm of cancer cells preferentially to the TCA cycle (Warburg effect) (29). Furthermore, elevated glucose stimulates glycolysis and inhibits aerobic respiration in tumors (Crabtree effect) (30). Thus, high-glucose BMs promote conversion of glucose to pyruvate and NADH generation but inhibit production of substrates for DNFA via the TCA cycle (31, 32). In general, the glucose content of DMEM may distort normal cellular behavior and thus render it unsuitable for molecular study of DNFA transcription regulation. Both aerobic respiration and anaerobic glycolysis contribute to glucose catabolism in cancer cells cultured with RPMI-1640 medium (33). Therefore, between the two common BMs, RPMI-1640 seems preferable for lipogenesis-related investigations of cancer cells.

The commercially available insulin, transferrin, and selenium (ITS) supplement is a serum replacement, supporting cell survival and growth but containing no lipids. ITS supplement has been used for *in vitro* culture of mesenchymal stem cells isolated from adipose or cartilage tissues, to maintain their differentiation and proliferation capacities for tissue transplantation (34, 35). Insulin is a growth factor with a mitogenic effect in cell culture that promotes the uptake of glucose and amino acids (36, 37). In livers, insulin stimulates *SREBF1c* mRNA expression (22) as well as proteolytic maturation of the nuclear SREBP1c protein to stimulate DNFA gene expression (22, 38). Transferrin is a glycoprotein that transports Fe^3+^ in blood plasma and delivers Fe^3+^ to cells through binding to transferrin receptor on cell surface (39). Fe^3+^ is an essential component of heme-containing enzymes like cytochromes for oxidative phosphorylation process and various non-heme iron enzymes, such as ribonucleotide reductase for DNA synthesis (40). Transferrin provides Fe^3+^ necessary to support cell survival and proliferation in culture (41). Selenium is required for proper function of antioxidant enzymes, including glutathione peroxidase and thioredoxin reductase, in which selenocysteine is indispensable for their catalytic activities (42). Components of ITS have been individually assembled in various serum-free media to support cell growth in cancer studies (43), but reports relevant to lipid metabolism are missing. Here, we introduce the combination of RPMI-1640 and ITS (ITS medium) as a straightforward serum-free medium condition for activating DNFA gene expression in melanoma cancer cell culture. The ITS medium stimulates cell growth, exhibits consistent effect across batches, and is free of confounding lipid factors. We expect that it may be adopted for investigations in lipid metabolism in cell lines from different cancer types and should facilitate *in vitro* screening for DNFA pathway inhibitors.

## Materials and Methods

### Cell culture reagent

The melanoma cell lines were obtained from the MGH Center for Molecular Therapeutics. Four melanoma cell lines from different stages of melanoma progression were used for these studies. MEL-JUSO is derived from a primary cutaneous melanoma (pre-metastatic) tumor (44, 45), WM1552C cell line is derived from a primary melanoma tumor with vertical growth phase in the patient (46, 47), HT-144 is a malignant melanoma cell line derived from a metastatic site (48, 49), and A375 is derived from a malignant melanoma tumor with highly metastatic amelanotic feature (50, 51). Cell lines were cultured in RPMI-1640 medium (21870092, Thermo Fisher Scientific) with 10% fetal bovine serum (Gibco), 2 mM L-glutamine (Gibco) and 50 U/ml penicillin-streptomycin (Gibco) in a 37 °C incubator with 5% CO_2_. The 0% FBS medium contained RPMI-1640 medium, with 2 mM L-glutamine (Gibco) and 50 U/ml penicillin-streptomycin (Gibco). The 1% ITS medium contained the RPMI-1640 medium with 1 × Insulin-Transferrin-Selenium (ITS-G, Thermo Fisher Scientific), 2 mM L-glutamine (Gibco) and 50 U/ml penicillin-streptomycin (Gibco).

#### Long-term culture in 1% ITS medium

HT-144, A375, and WM1552C cells were maintained in 25 cm^2^ flasks in 1% ITS medium. When melanoma cells were approximately 80% confluent, they were rinsed with sterile PBS solution and then detached with 0.5 mL trypsin solution (0.25%). The equivalent of 5 volumes of pre-warmed plating medium (mixture of 1% ITS and 10% FBS media in 9: 1 ratio) was added in order to inactivate trypsin. Cell suspension was pipetted into new flasks at 1:6 split ratio and then topped up with plating medium. Newly passaged cells were left at 37°C for 1.5 hours to recover and settle on flask. Then the plating medium was removed and changed to 1% ITS medium to remove any residual trypsin and FBS for continuous culture. HT-144 and A375 cells were passaged in 1% ITS medium every 5-7 days, whereas WM1552C cells were passaged in 1% ITS medium approximately every 10 days.

### Cell proliferation assays

#### AlamarBlue assay

Cells were seeded in 24-well plates (Corning) in RPMI-1640 medium with 10% FBS at a density of 2,000 cells per well for HT-144, MEL-JUSO and WM1552C, and 1,000 cells per well for A375, Sixteen hours after seeding, cells were washed twice with PBS buffer and then cultured in three different medium conditions for 6 days. Relative cell viability was quantified for cellular metabolic activities by alamarBlue cell viability reagent (DAL1025, Thermo Fisher Scientific). Fluorescence emission of reduced resorufin was measured at 590 nm wavelength using an Envision 2103 multilabel microplate reader (Perkin Elmer). Each data point represents average measurement from four replicate samples with duplicate measurement. Because the culture medium influences resazurin fluorescence, the background fluorescence of medium-only with alamarBlue reagent was subtracted for data normalization.

#### CyQuant assay

Cells were seeded in 96-well plates (Corning, 3610) in RPMI-1640 medium with 10% FBS at a density of 1,000 cells per well for HT-144, MEL-JUSO and WM1552, and 500 cells per well for A375. Sixteen hours after seeding, cells were then cultured in three different medium conditions for six days. Relative live cell numbers were quantified based on DNA content and membrane integrity with CyQuant direct cell proliferation assay (C35011, Thermo Fisher Scientific). Green fluorescent nucleic acid stain signal was measured with FITC filter set on the Envision 2103 multilabel microplate reader (Perkin Elmer). The background fluorescence of medium-only with CyQuant reagent was subtracted for data normalization.

### Reverse transcription quantitative PCR (RT-qPCR)

Total RNA was isolated from cultured cells using the RNeasy mini kit (Qiagen) and treated with RNase-free DNase (Qiagen). RNA concentrations were quantified with Qubit™ RNA BR assay kit (Thermo Fisher Scientific). One μg RNA was used for cDNA synthesis with RNA to cDNA EcoDry™ premix (TaKaRa) containing both random hexamer and oligo(dT)18 primers (Double Primed). qPCR was carried out in triplicates on a LightCycler^®^ 480 instrument (Roche) using LightCycler^®^ 480 SYBR green I master (Roche). qPCR primers were pre-designed by MGH primer bank (https://pga.mgh.harvard.edu/primerbank/) and the primer sequences are listed in Table 1. Relative gene expression levels were calculated using the 2^−ΔΔCt^ method, normalized to the 18S housekeeping gene, and the mean of negative control samples was set to 1.

**Table 1.**
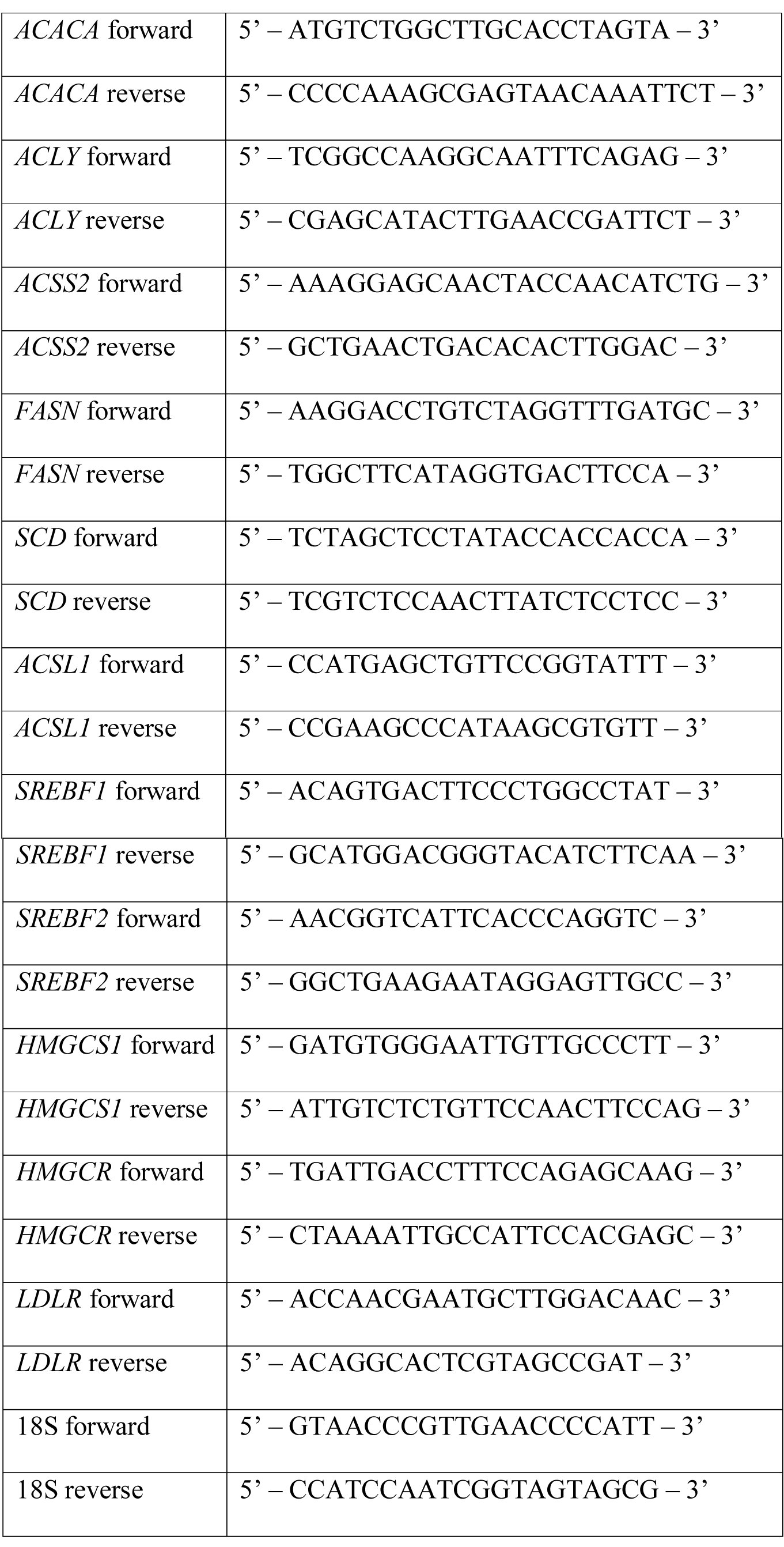
Primers used for RT-qPCR.siRNA transfection

### siRNA transfection

The human-specific siRNAs targeting *SREBF1* (6720) and *SREBF2* (6721) were pre-designed ON-TARGETplus SMARTpool siRNA reagents from Dharmacon. Each ON-TARGETplus SMARTpool siRNA was a mixture of four siRNA duplexes. siRNAs were suspended in RNase-free 1 × siRNA Buffer (Dharmacon) to yield 20 μM stock solutions. HT-144 cells were transfected with siRNAs at a final concentration of 50 nM using Lipofectamine RNAiMAX transfection reagent (Thermo Fisher Scientific) at a density of 2 × 10^5^ cells/well in a 6-well plate. For each transfection, 2.5 μl of siRNA stock solution was mixed with 4 μl of Lipofectamine RNAiMAX in 200 μl of Opti-MEM I medium and then incubated for 10-20 minutes at room temperature. HT-144 cells were diluted in 10% FBS/RPMI-1640 medium without antibiotics so that 800 μl medium contains 2.5 × 10^5^ HT-144 cells. 200 μl of siRNA/Lipofectamine RNAiMAX complexes was mixed with 800 μl of the diluted HT-144 cells in one well of a 6-well plate. Transfected HT-144 cells were incubated at 37°C for 16 hours before medium changes.

### Immunoblot assay

HT-144 cells were seeded in 10 cm^2^ plates in 10% FBS/RPMI-1640 medium at day one. Sixteen hours after seeding, cells were washed twice with PBS buffer and then cultured in different medium conditions. Total cell lysate was harvested with RIPA buffer containing protease inhibitors (protease inhibitor cocktail tablets, Roche) on the third day. Nuclear and cytoplasmic protein fractions were prepared with NEPER™ nuclear and cytoplasmic extraction reagents (Thermo Fisher Scientific). Protein samples were separated on the 4-15% Mini-PROTEAN^®^ TGX™ precast SDS-PAGE gels (Bio-Rad) and then transferred to polyvinyl difluoride (PVDF) membranes (Immobilon-P, Millipore) for immunoblot analysis. The following primary antibodies were used: mouse anti-SREBP1 (IgG-2A4, BD Biosciences), rabbit anti-FASN (C20G5, Cell Signaling), rabbit anti-SCD (23393-1-AP, Proteintech), rabbit anti-ACSL1 (D2H5, Cell Signaling), rabbit anti-histone H3 (9715, Cell Signaling) and rabbit anti-beta-actin (13E5, Cell Signaling). After being incubated with primary antibodies overnight in PBST solution with 5% non-fat dry milk, immunoblot membranes were probed with HRP-conjugated affinity-purified donkey anti-mouse or anti-rabbit IgG (GE Healthcare) as secondary antibodies and visualized with the immobilon Western Chemiluminescent HRP substrate (Millipore).

### Chromatin immunoprecipitation (ChIP) quantitative PCR (ChIP-qPCR)

For each ChIP assay, 5 × 10^7^ HT-144 cells were used. HT-144 cells were seeded in 10% FBS medium, and then medium was replaced with 10% FBS or 1% ITS medium and cultured for 24 hours before ChIP-qPCR analyses. Chromatin from HT-144 cells was fixed with 1% formaldehyde (Polysciences) and prepared with Magna ChIP™ HiSens chromatin immunoprecipitation kit (EMD Millipore). Nuclei were sonicated on a sonic dismembrator 550 (Fisher Scientific) with a microtip (model 419) from Misonix Inc. Lysates were sonicated on ice with 10 pulses of 20 seconds each (magnitude setting of 3.5) and a 40-sec rest interval. The supernatant was used for immunoprecipitation with the following antibodies: rabbit anti-SREBP1 (H-160, Santa Cruz Biotechnology), rabbit anti-CBP (C-22, Santa Cruz Biotechnology), rabbit anti-RNA polymerase II (8WG16, BioLegend) and rabbit anti-histone H3 (acetyl K27) (ab4729, Abcam). qPCR reactions in triplicates were performed on a LightCycler^®^ 480 instrument (Roche) using LightCycler^®^ 480 SYBR green I master (Roche). The ChIP-qPCR primers were designed with software Primer 3. The primer sequences are listed in Table 2.

**Table 2.**
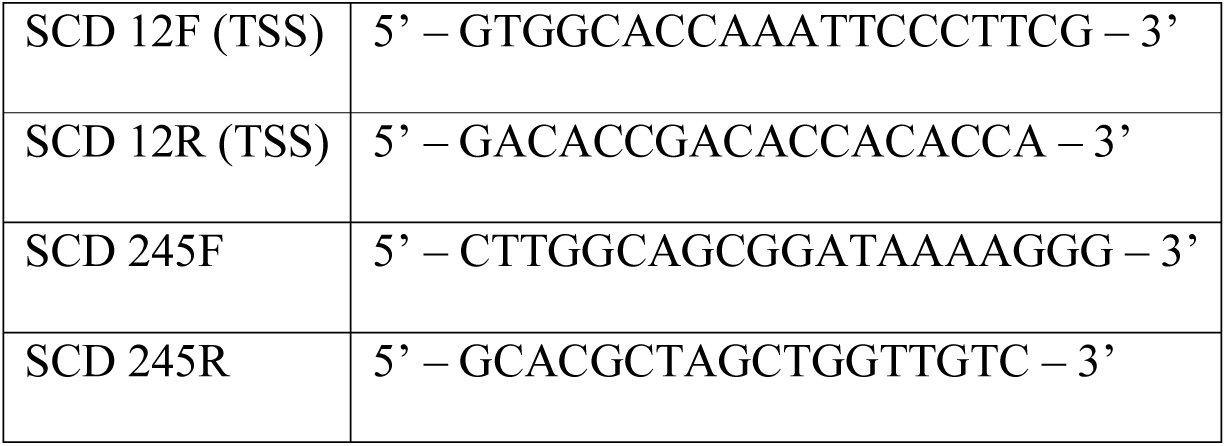
Primers used for ChIP-qPCR.

## Results

### A lipid-free and insulin-supplemented (ITS) medium supports melanoma cell proliferation

To assess the utility of the 1% ITS medium for culturing melanoma cell lines and evaluating DNFA gene expression, we first performed time-course analysis to measure growth rates of melanoma cell lines under three different cell culture medium conditions: RPMI-1640 supplemented with either 10% FBS, 0% FBS, or 1% ITS. Four different melanoma cell lines were tested: HT-144, A375, WM1552C and MEL-JUSO. HT-144 and A375 were derived from human metastatic melanomas (48, 51), whereas WM1552C was derived from a stage III superficial spreading primary melanoma (47), and MEL-JUSO was derived from a primary human melanoma (52). Viable melanoma cells were quantified using alamarBlue daily for six days. We found that HT-144, A375 and WM1552C cells cultured in 10% FBS and 1% ITS medium conditions displayed a time-dependent increase of fluorescence reads in the alamarBlue assay (Fig 1A, S1A - S1B Fig). We also observed increased cell numbers with light microscopy, confirming that the cell lines proliferated in both 10% FBS and 1% ITS medium conditions. We observed a growth plateau of HT-144, A375 and WM1552C cells in 0% FBS medium, which lacks growth factors such as insulin, and in which cells therefore remain quiescent. MEL-JUSO cells did not display a time-dependent increase of fluorescence reads in 0% FBS and 1% ITS medium conditions with alamarBlue assay (S1C Fig), which indicates that MEL-JUSO cells did not proliferate in these two medium conditions, possibly related to its less transformed phenotype (52).

**Fig 1.**
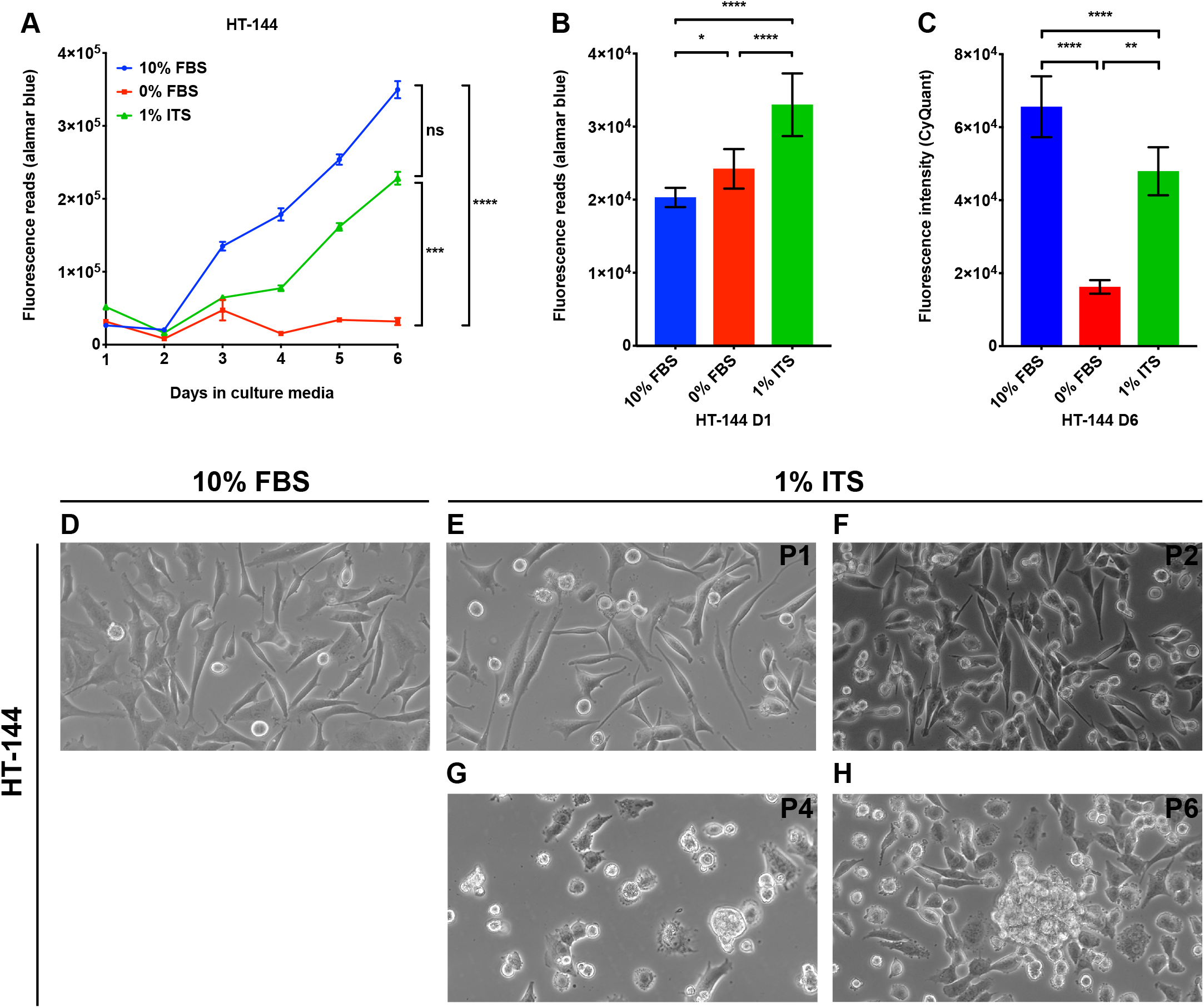
A lipid-free and insulin-supplemented medium supports proliferation of HT-144 melanoma cells. (A) HT-144 cells were seeded in 10% FBS medium at day zero. On day one, cells were washed with PBS and changed to the indicated medium conditions. Cell proliferation was measured with alamarBlue assay daily for six days. Each data point represents the mean ±SD of quadruplicate samples. The results were analyzed using two-way repeated measures ANOVA followed by *post hoc* Tukey’s multiple comparison tests. For culture time, F = 1505, P < 0.0001; for culture condition, F = 2153,P < 0.0001; for interaction between culture time and condition, F = 409.4, P < 0.0001. (B) HT-144 cells were seeded in 10% FBS medium at day zero. On day one, cells were washed with PBS and changed to the indicated medium conditions. alamarBlue assay was performed on the cells cultured in the indicated medium for one hour. Results were analyzed using one-way ANOVA followed by *post hoc* Tukey’s multiple comparison tests. F = 202.7. Significant differences between medium conditions are indicated as *P < 0.05, **P < 0.01, ***P < 0.001 and ****P < 0.0001. ns, not significant. (C) HT-144 cells were seeded in 10% FBS medium at day zero. On day one, cells were washed with PBS and changed to the indicated medium conditions. CyQuant assays were performed on the cells cultured in the indicated medium at day six. Each data bar represents average measurement of five replicate samples. Results were analyzed using one-way ANOVA followed by *post hoc* Tukey’s multiple comparison tests. F = 80.77, P < 0.0001. Significant differences between medium conditions are indicated as *P < 0.05, **P < 0.01, ***P < 0.001 and ****P < 0.0001. ns, not significant. (D - H) HT-144 cells were cultured in RPMI medium with 10% FBS or 1% ITS supplement for multiple passages. Morphologies of cells cultured in 1% ITS medium from passage one (P1) to passage six (P6) were monitored by light microscopy with 40 × objective and 10 × ocular lens.

The alamarBlue assay measures the fluorescence emission of resorufin molecules converted from resazurin by reductants such as NADPH or NADH (53, 54). The alamarBlue assay is thus an indicator of cellular NADPH/NADH concentration in living cells. Because cytosolic NADPH serves as the reductant for biosynthesis pathways such as lipid and nucleic acid synthesis, alamarBlue assay may provide a hint regarding DNFA activity. Therefore, we compared the alamarBlue reads of cells cultured in different medium conditions at day one, when they had the same cell numbers in the three medium conditions. HT-144 cells cultured in 1% ITS medium had the highest fluorescence reads, whereas those cultured in 10% FBS medium had the lowest reads (Fig 1B). This result suggests that even when the cell number remains the same, HT-144 cells cultured in 1% ITS possibly have higher NADPH/NADH concentrations. We observed similar increases of alamarBlue reads when culturing A375 and WM1552C cells in 1% ITS medium condition (S1D – S1E Fig), and an insignificantly small increase in MEL-JUSO cells cultured in 1% ITS medium compared to 10% FBS medium (S1F Fig).

To ensure that metabolic activities measured by the alamarBlue assay represent the cell proliferation rate (55), we used CyQuant, a DNA-content based assay, for confirmation. We measured the DNA content of four melanoma cell lines cultured in three different medium conditions after six days of culture. We observed that DNA contents for HT-144, A375 and WM1552C cell lines were significantly higher in 1% ITS medium than those in 0% FBS medium (Fig 1C, S1G and S1H Fig). We observed no difference in DNA content in MEL-JUSO between 1% ITS and 0% FBS medium conditions (S1I Fig), consistent with the cell proliferation results from alamarBlue assays (S1C Fig). CyQuant assay results validated the finding that A375, HT-144 and WM1552C cells proliferate in lipid-free and insulin-supplemented medium. Since 1% ITS medium does not contain any external lipids, we reasoned that the lipids employed under this condition for membrane synthesis and other cellular needs during proliferation of A375, HT-144 and WM1552C cells are entirely derived from DNFA.

To account for the possibility that elevated DNFA acts as short-term adaptation to support cell proliferation after transfer to 1% ITS medium for culture, we investigated the four melanoma cell lines in that medium for long-term cell growth. We observed that cell lines derived from metastatic melanomas, A375 and HT-144, are able to continuously proliferate under 1% ITS medium condition and were sub-cultured by six passages over 6 weeks (Fig 1E – 1H, S2B – S2E Fig). WM1552C, a primary melanoma cell line, proliferated much more slowly in 1% ITS medium than in 10% FBS medium, but we were able to subculture this line by four passages over 6 weeks (S2G – S2J Figs). We also noticed morphological changes of A375, HT-144 and WM1552C cells under long-term 1% ITS medium culture. The cells changed shape from an adherent and flattened appearance in 10% FBS (Fig 1D, S2A and S2F Fig) to a rounded form at late passages in 1% ITS (Fig 1H, S2E and S2J Fig). HT-144 and particularly A375 cells formed small spheroids, with some cells floating as single cells. MEL-JUSO, a primary melanoma cell line, failed to proliferate or survive passage in 1% ITS medium (S2K – S2M Fig). These results suggest that the ability to proliferate in 1% ITS medium condition varies among melanoma cell lines, largely in line with degree of progression from primary to metastatic cell state.

### ITS medium increases DNFA and DNCS gene expression in melanoma cells

To understand whether proliferation and cell survival in 1% ITS medium were linked to elevated DNFA activity, we performed RT-qPCR analyses of DNFA mRNA, including *ACLY, ACSS2, ACACA, FASN, SCD*, and *ACSL1*. We also examined the expression of *de novo* cholesterol synthesis (DNCS) genes, including *HMGCS1* and *HMGCR*. HT-144 cells were seeded in 10% FBS medium, and then medium was replaced with 0% FBS or 1% ITS medium and cultured for 24 hours before RT-qPCR analyses. We observed a significant increase in the expression of DNFA (Fig 2A-2F), DNCS (Fig 2G and 2H) and *LDLR* (Fig. 2I) genes when comparing cells cultured in 0% FBS and 1% ITS medium conditions to cells in 10% FBS medium. We observed lower DNFA and DNCS gene expression in cells cultured in 10% FBS medium than in 0% FBS medium, consistent with repression of DNFA and DNCS by lipids derived from the serum supplement.

**Fig 2.**
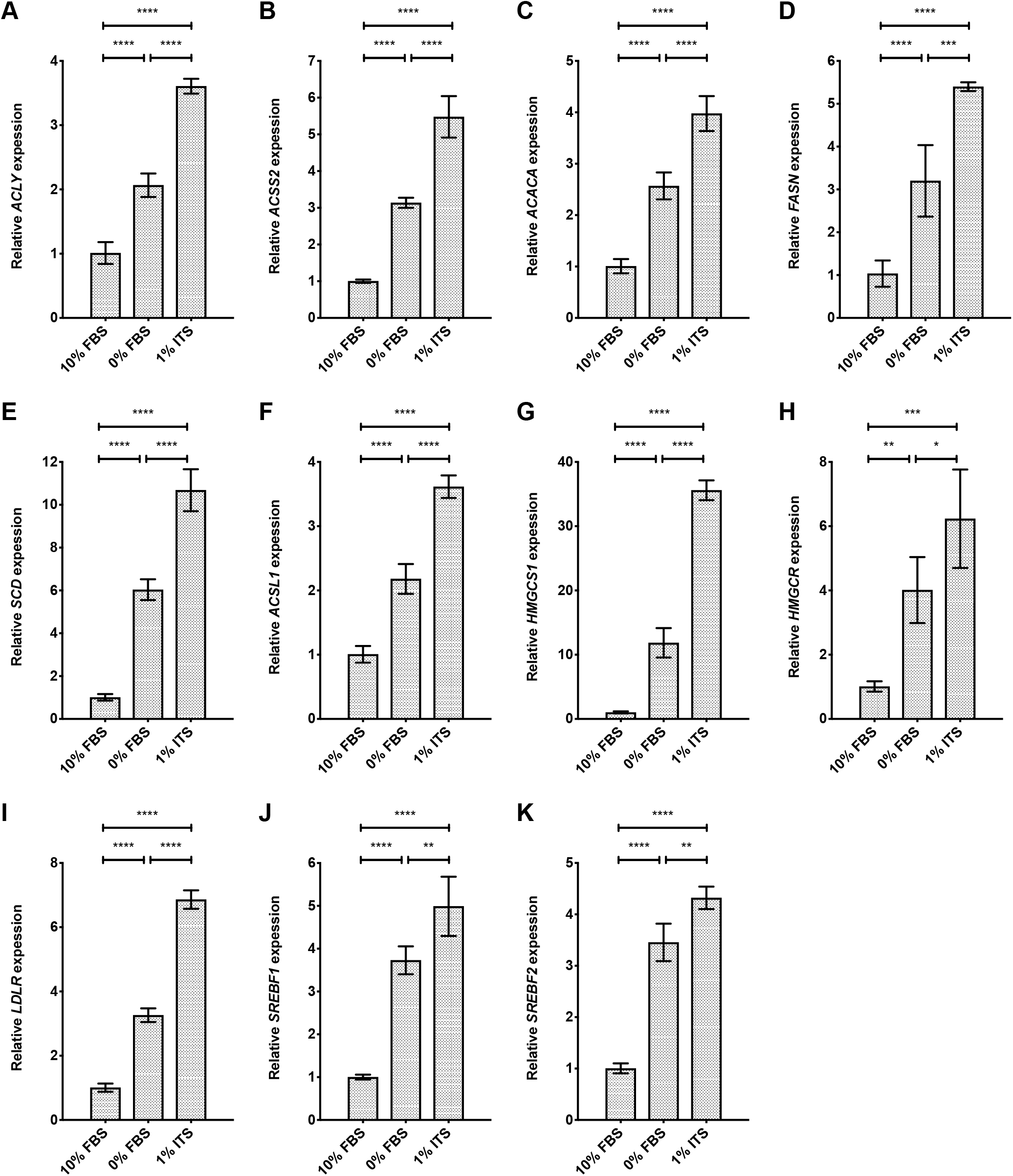
Lipid depletion combined with insulin supplement yields increased expression of lipogenic genes in HT-144 cells. (A-K) The expression level of DNFA and DNCS genes was analyzed by RT-qPCR assay. HT-144 cells were cultured in 10%, 0% FBS or 1% ITS medium for 24 hours. Significant differences between medium conditions are indicated as *P < 0.05, **P < 0.01, ***P < 0.001 and ****P < 0.0001 using one-way ANOVA followed by *post hoc* Tukey’s multiple comparison tests. ns, not significant. Each data point represents the mean ±SD of results from quadruplicate samples.

Fig 1 indicates that HT-144 cells remain quiescent in 0% FBS medium and proliferate in 1% ITS. Quiescent HT-144 cells have enhanced DNFA and DNCS gene expression, even though there is no requirement for membrane lipid synthesis to support proliferation, and the reasons for this remain unclear. DNFA and DNCS gene expression is elevated in 1% ITS medium as compared with 0% FBS medium, which may suggest that the insulin component of the ITS supplement contributes to stimulation of DNFA gene expression.

We further observed significantly increased expression of *SREBF1* and *SREBF2* genes in cells cultured in 0% FBS and 1% ITS medium conditions, compared to 10% FBS medium (Fig 2J and 2K). These results suggest that SREBP1 and SREBP2 are transcriptionally activated in lipid-free and insulin-supplemented conditions. Because SREBP1/2 can bind to sterol regulatory elements (SRE) located in their own promoters, SREBP1/2 proteins possibly regulate their mRNA production through auto-activation (56) and this mechanism may also contribute to elevated DNFA and DNCS gene expression.

We additionally performed gene expression analyses of DNFA and DNCS pathways in MEL-JUSO cells, which failed to proliferate in 1% ITS medium. MEL-JUSO cells were seeded in 10% FBS medium, and then medium was replaced with 0% FBS or 1% ITS medium and cultured for 24 hours before RT-qPCR analyses. We observed a significant increase in the expression of DNFA (S3A-S3F Fig), DNCS (S3G, S3H Fig) and *LDLR* (S3I Fig) genes when comparing cells cultured in 0% FBS and 1% ITS medium conditions to cells cultured in 10% FBS medium. However, the increase in expression of *SREBF1* (Fig 2E and S3E Fig) and certain DNFA genes such as *ACACA* (Fig 2C and S3C Fig) and *SCD* (Fig 2J and S3J Fig) was much lower in MEL-JUSO cells than in HT-144 cells, when comparing 1% ITS medium conditions to 10% FBS medium. These results suggest that DNFA activities can be elevated as a short-term response to lipid-free condition and insulin supplement in cells that cannot proliferate by relying upon DNFA as a sole source of lipids. We reason that the robustness and sustainability of DNFA elevation in melanoma cells probably influences proliferative capacity.

### ITS medium increases DNFA and DNCS gene expression *via* SREBP1 and SREBP2, respectively

SREBP1 and SREBP2 are master transcription regulators of DNFA and DNCS pathways, respectively (57). To verify cellular dependence on SREBP1 and SREBP2 for lipid biosynthesis pathways, we transfected HT-144 cells with pooled siRNAs to deplete mRNAs encoding *SREBF1* and *SREBF2*. The transfected cells were cultured in 10% FBS, 0% FBS or 1% ITS medium conditions for two days and then assayed with RT-qPCR for DNFA (Fig 3) and DNCS (Fig 4) gene expression.

**Fig 3.**
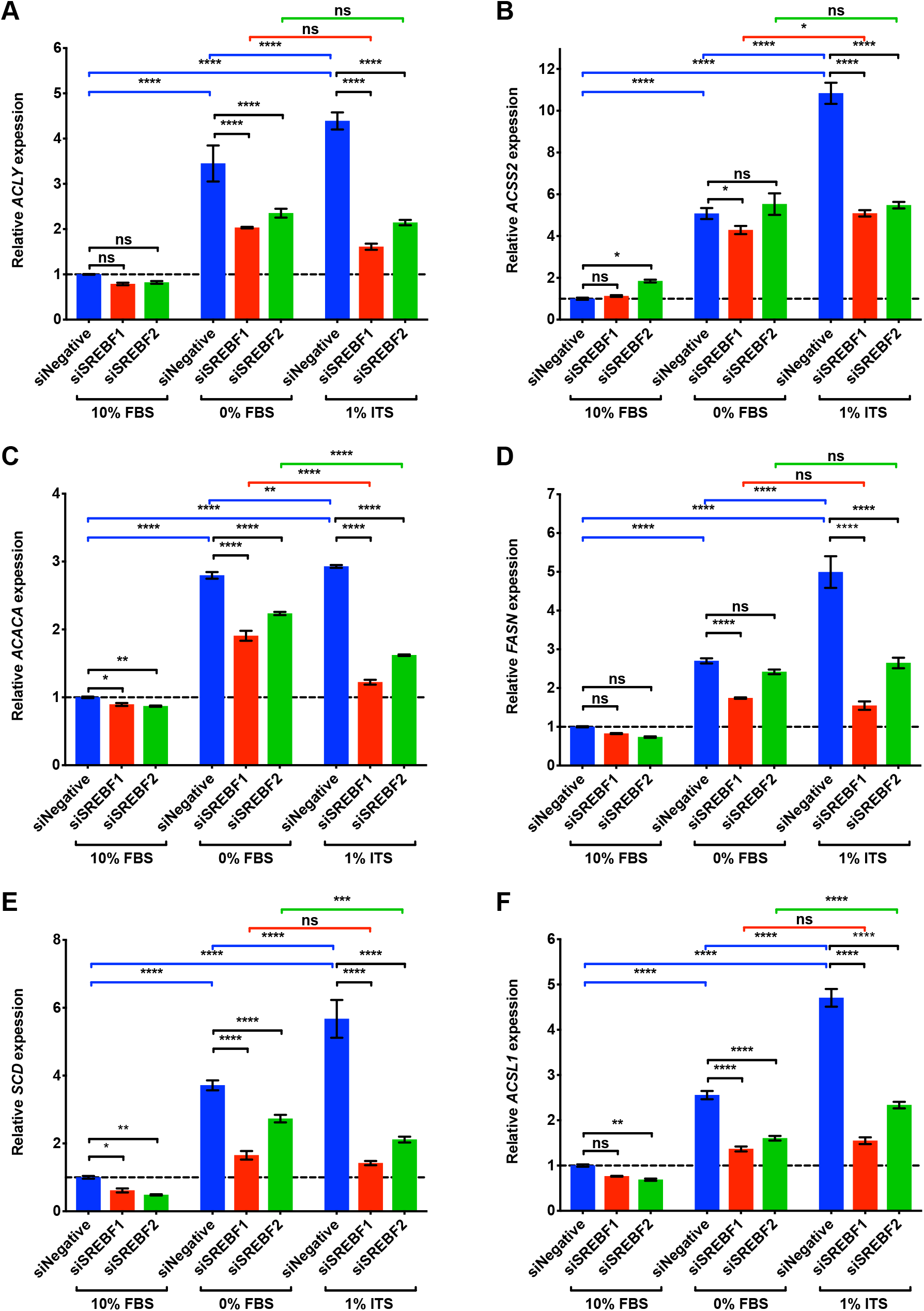
DNFA gene expression of HT-144 cells increases in ITS medium, and this increase is dependent upon SREBP1. (A-F) HT-144 cells were transfected over one day with siRNAs at 50 nM concentrations for non-target control, *SREBF1* or *SREBF2*. Transfected cells were cultured in 10%, 0% FBS or 1% ITS for two more days before RT-qPCR analyses. RT-qPCR results are presented as expression of DNFA genes relative to their expression under scramble siRNA treatment (siNegative) in 10% FBS medium (set as 1 and marked with a dashed line). When a significant interaction between siRNA treatment and medium condition was detected by two-way ANOVA, individual gene expression was compared within groups by *post hoc* Tukey’s multiple comparison tests. Each data point represents the mean ±SD of triplicate samples. *, P < 0.05; **, P < 0.01; ***, P < 0.001; ****, P < 0.0001.

**Fig 4.**
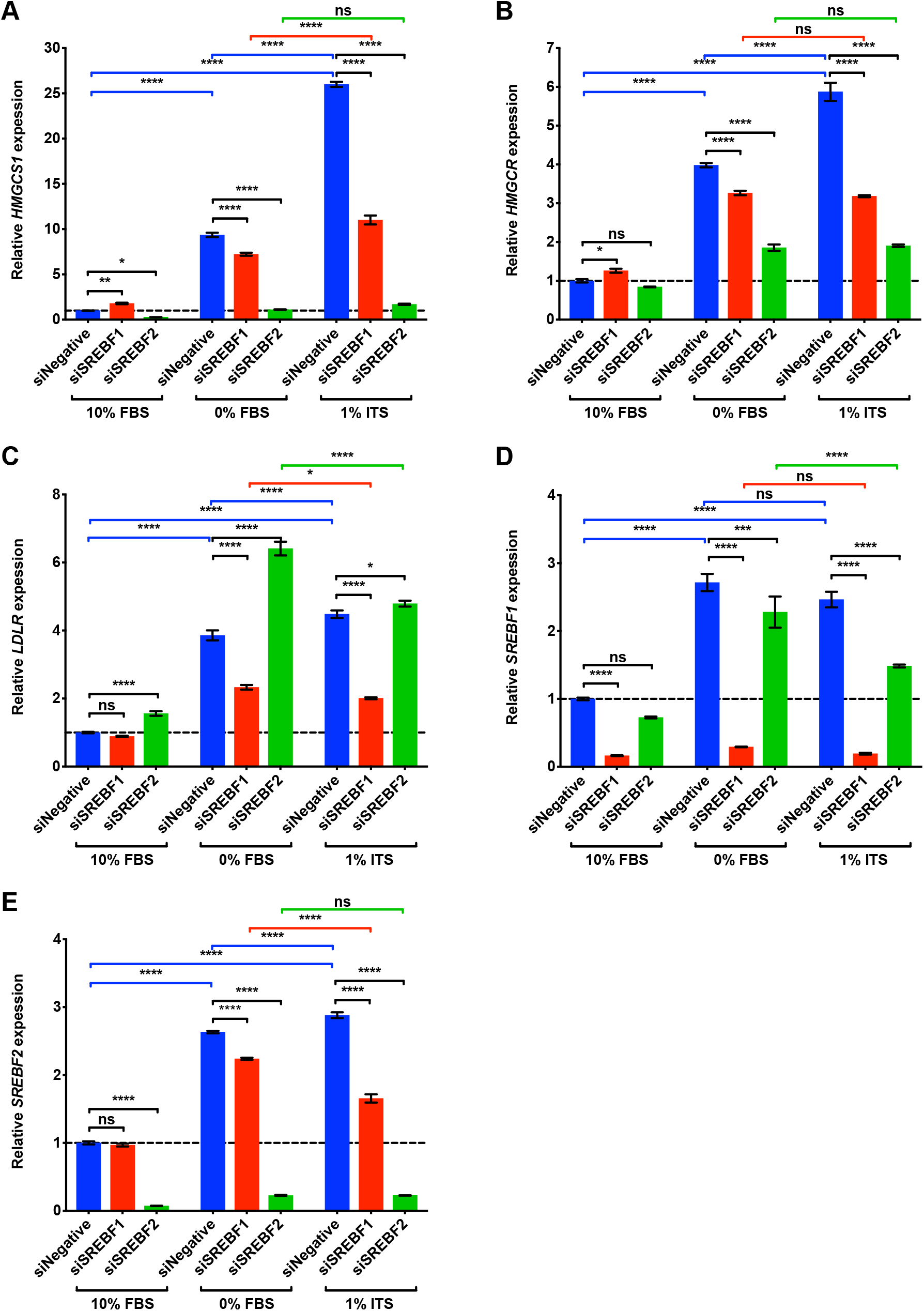
DNCS gene expression of HT-144 cells increases in ITS medium, and this increase is dependent upon SREBP2. (A-E) HT-144 cells were transfected over one day with siRNAs at 50 nM concentrations for non-target control, *SREBF1* or *SREBF2*. Transfected cells were cultured in 10%, 0% FBS or 1% ITS for two more days before RT-qPCR analyses. RT-qPCR results are presented as expression of genes relative to their expression under scramble siRNA treatment (siNegative) in 10% FBS medium (set as 1 and marked with a dashed line). When a significant interaction between siRNA treatment and medium condition was detected by two-way ANOVA, individual gene expression was compared within groups by *post hoc* Tukey’s multiple comparison tests. Each data point represents the mean ±SD of triplicate samples. *, P < 0.05; **, P < 0.01; ***, P < 0.001; ****, P < 0.0001.

In the group treated with scrambled siRNA (blue bars in Figs 3 and 4), we found that DNFA and DNCS gene expression was significantly elevated in 0% FBS and 1% ITS medium conditions compared to 10% FBS, consistent with our observations in Fig 2. However, DNFA gene expression is significantly lower in the *SREBF1* depleted group than in the scramble siRNA group, particularly in 0% FBS and 1% ITS medium conditions (Fig 3A-3F, statistical comparisons marked in black). DNCS gene expression is significantly lower in the *SREBF2* depleted group than in the scramble siRNA group, in 0% FBS and 1% ITS medium conditions (Fig 4A and 4B, statistical comparisons marked in black). Two-way ANOVA analysis detects a significant interaction between *SREBF1/2* depletion and medium condition for DNFA and DNCS gene expression. This result provides a statistical indication that SREBP1 and SREBP2 participate in activation of lipogenic gene expression in 0% FBS and 1% ITS conditions. We further found that *SREBF1* siRNA has the greatest effect on DNFA gene expression (Fig 3A-3F), and *SREBF2* siRNA has the greatest effect on DNCS gene expression (Fig 4A and 4B). *LDLR* appears to be regulated primarily through SREBP1 rather than SREBP2 in HT-144 cells (Fig 4C). Our results are consistent with known roles for SREBP1 and SREBP2 in activation of DNFA and DNCS gene expression (57).

In the scrambled siRNA treated groups, we observed significantly higher DNFA and DNCS gene expression in 1% ITS than in 0% FBS medium, suggesting additional influence of the ITS supplement on DNFA gene expression (Figs 3 and 4, statistical comparisons marked in blue). In the *SREBF1*-depleted group, we observed no significant increase of DNFA gene expression when comparing 1% ITS to 0% FBS condition (Fig 3A-3F, statistical comparisons marked in red). Similarly, in SREBF2-depleted group there is no significant elevated expression of DNCS genes, *HMGCS1* and *HMGCR*, in 1% ITS medium compared to 0 % FBS (Fig 4A and 4B, statistical comparisons marked in green). Altogether, these results support the utility of the 1% ITS medium condition to study SREBP1- and SREBP2-targeted interventions to inhibit DNFA and DNCS respectively, in the absence of confounding serum lipids.

### ITS medium increases the level of nuclear SREBP1

To confirm that lipid-free and/or insulin-supplemented medium conditions influence SREBP1 protein production in general and its nuclear form in particular, we examined the cytoplasmic and nuclear levels of SREBP1 in HT-144 cells cultured under the three medium conditions (Fig 5A). Our results revealed a dramatic increase of nuclear SREBP1 in 0% FBS and 1% ITS medium conditions. However, we did not observe a decrease of full-length SREBP1 in the cytoplasmic fraction from 0% FBS and 1% ITS medium conditions. The overall SREBP1 protein level increased in 0% FBS and 1% ITS medium conditions, which is likely due to transcriptional activation of the *SREBF1* gene (Fig 2J).

**Fig 5.**
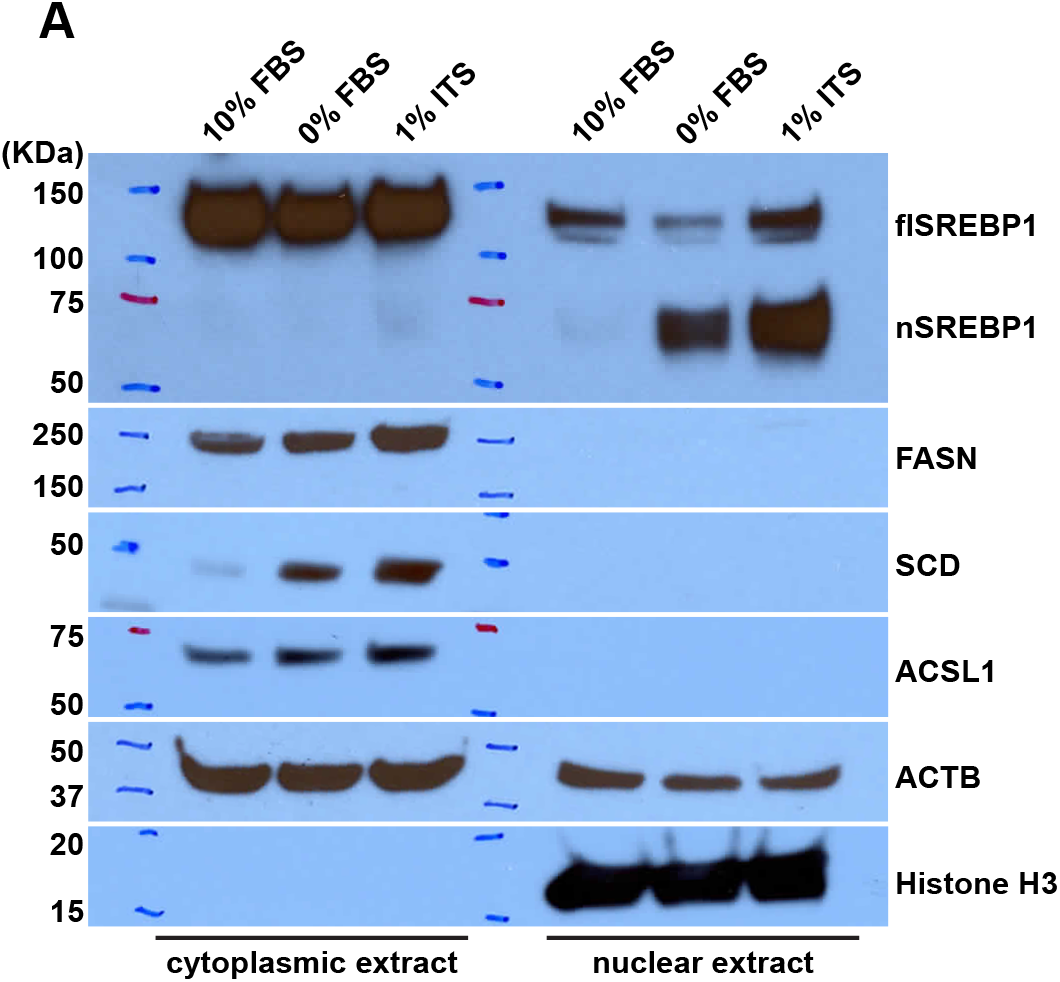
Lipid depletion combined with insulin supplement yields increased expression of nuclear SREBP1. (A) The cytoplasmic and nuclear fractions of HT-144 cells cultured in 10%, 0% FBS or 1% ITS medium were isolated for Western blot analysis. HT-144 cells in 1% ITS medium have increased levels of nuclear SREBP1 protein and lipogenic enzymes, detected with the indicated antibodies. Histone H3 serves as the positive control for nuclear fractionation during biochemical preparation of the cell samples.

Consistent with the increased mRNA for DNFA genes (Fig 2D-2F), there is increased production of DNFA enzymes including FASN, SCD and ACSL1 in the cytoplasmic fraction from 0% FBS and 1% ITS medium conditions. The increase of DNFA enzyme production correlates with the increase of nuclear SREBP1. We interpret this as evidence that lipid depletion combined with insulin supplement greatly enhances the abundance of the SREBP1 nuclear form and thus stimulates DNFA enzyme production.

### Culture in ITS medium increases SREBP1 binding at the *SCD* gene promoter

Finally, we investigated medium condition impact upon the molecular mechanism by which nuclear SREBP1 controls DNFA gene expression. We used chromatin immunoprecipitation (ChIP)-qPCR to examine the occupancy of SREBP1, transcriptional co-activator CBP (58, 59), RPB1 (the largest subunit of RNA polymerase II) and histone marker H3K27Ac (a marker for active enhancers and promoters) (60) on the promoter of DNFA gene *SCD* (Fig 6A-6D). We observed a significant increase of SREBP1 and CBP binding at the *SCD* transcription start site (TSS) in 1% ITS relative to 10% FBS (Fig 6A and 6C, statistical comparisons marked in black). A corresponding slight increase of RBP1 at the *SCD* TSS site is not statistically significant, however we observe a modest but significant increase in RPB1 occupancy downstream of TSS in the gene body (Fig 6B, statistical comparisons marked in black), consistent with active RNA polymerase II elongation (61). 1% ITS does not significantly increase the H3K27Ac signal at the *SCD* TSS nor downstream in the gene body compared to 10% FBS (Fig 6D, statistical comparisons marked in black). However, we observed abundant H3K27Ac signals in the gene body for both 10% and 1% ITS (Fig 6D, statistical comparisons marked in blue and red). This result suggests that the *SCD* gene body may be constitutively active, with open chromatin architecture.

**Fig 6.**
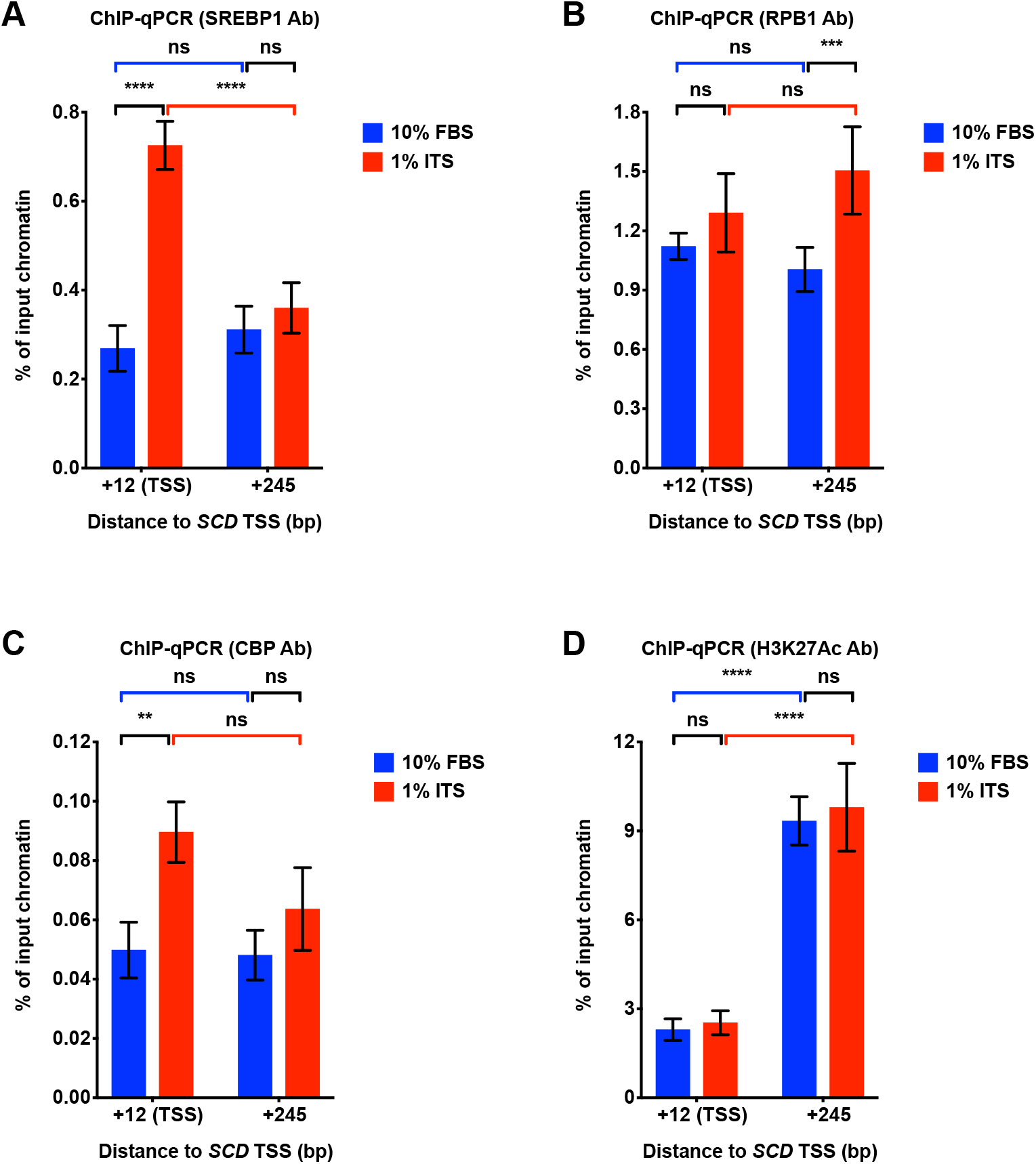
Lipid depletion combined with insulin supplement increased SREBP1 binding at the *SCD* gene promoter. (A-D) ChIP-qPCR analyses for enrichment of transcriptional regulation factors on the TSS and gene body region (245 bp downstream of TSS) of the *SCD* gene locus. qPCR was used to quantify chromatin immunoprecipitated with the indicated antibodies from HT-144 cells cultured in 10% FBS or 1% ITS medium. Quantification of enrichment was determined as percentage of input chromatin before immunoprecipitation. Each data point represents the mean ±SD of triplicate samples. Two-way ANOVA was used to identify any significant interaction between gene locus and medium condition on influencing binding of indicated factors in ChIP assay. (A) SREBP1 binding. For interaction between gene locus and medium, F = 86.24, P < 0.0001; (B) RNA polymerase II binding. F = 6.245, P = 0.0213. (C) CBP binding. F = 3.848, P = 0.0854. (D) H3K27Ac binding. F = 0.09706, P = 0.7586. Significant differences of binding are indicated as *P < 0.05, **P < 0.01, ***P < 0.001 and ****P < 0.0001 using two-way ANOVA followed by *post hoc* Tukey’s multiple comparison tests. ns, not significant.

Two-way ANOVA analysis detects significant interactions between gene locus and medium condition in SREBP1 and RNA polymerase II ChIP-qPCR data. This result indicates that 1% ITS significantly promotes enrichment of SREBP1 at TSS region and RNA polymerase II at gene body region. We reason that, in 1% ITS medium, the likely transcription regulation mechanism after SREBP1 binding involves promotion of the transition from RNA polymerase II pausing to active elongation (61).

## Discussion

We report here that a serum-free and insulin-supplemented culture medium condition, 1% ITS supplemented RPMI-1640, supports DNFA and DNCS pathway activation as well as proliferation and survival of human melanoma cell lines. Under this condition, HT-144 cells proliferate while relying entirely on *de novo* lipid synthesis to meet lipid requirements. Expression of DNFA and DNCS enzymes increases significantly in cells when cultured in 1% ITS as compared with other cell culture conditions. We found that 1% ITS medium activates DNFA and DNCS gene expression through the transcription regulators SREBP1 and SREBP2, respectively. In particular, culturing cells in 1% ITS medium promoted the transcription activation of SREBP1 and accumulation of nuclear SREBP1 protein, as compared with other medium conditions, and cells cultured in 1% ITS medium exhibited further increased binding of SREBP1 at a DNFA gene promoter, consistent with high SREBP1 - dependent gene activation under this medium condition.

Unlike free fatty acids, free cholesterol and cholesteryl esters are only minimally soluble in blood and must be transported within lipoproteins (62). Lipoprotein-deficient serum (LPDS) is thus more efficient in removing cholesterol and cholesteryl esters than fatty acids. LPDS has been used in lieu of full serum as an activating medium for SREBPs, because it alleviates sterol inhibition (63). However, LPDS contains free fatty acids that enter cells through passive membrane diffusion or active transport through membrane receptors (64). LPDS has therefore also been used for studies of cellular fatty acid uptake and lipid storage from extracellular fatty acids (65). By contrast, ITS medium contains no external free fatty acids or lipoproteins. Membrane lipids for cell proliferation in ITS medium rely exclusively upon DNFA and DNCS. Thus, ITS represents an attractive culture medium as compared with LPDS for studying DNFA-supported proliferation and cell survival of cancer cells.

Our results are consistent with the notion that active DNFA is sufficient for cell proliferation in some malignant melanoma cell lines but not in others. Lipid depletion in 0% FBS activated DNFA and DNCS gene expression, possibly due to removal of exogenous cholesterol, the classic feedback inhibitor of SREBP processing and lipid synthesis (66). HT-144 cells are able to persist and survive under these conditions, but become quiescent, likely due to removal of growth factors in the serum necessary to support active proliferation. HT-144 cells proliferate, still with elevated DNFA and DNCS expression, in the presence of insulin, the only growth factor provided by 1% ITS. Similarly, DNFA gene expression was upregulated in MEL-JUSO cells when cultured in 1% ITS medium, even though MEL-JUSO failed to proliferate in this condition. This data supports the model where insulin present in the serum free ITS medium has both metabolic and mitogenic functions in cancer cells, promoting (with varying efficacy) cell survival and active proliferation. The entire regulatory regime for DNFA and its role in cell survival, in which insulin participates when present, is not yet fully clear but may involve the AKT/GSK3 signaling pathway (67, 68). Interestingly, two metastatic melanoma cell lines seem able to proliferate using only DNFA-derived lipids, whereas the two primary melanoma cell lines evaluated have either limited or no proliferative capacity when serum lipids are absent. Our study of four cell lines is too small to yield a conclusion on DNFA in metastatic vs primary tumors, but at least hints that DNFA and DNCS are important to tumor malignancy. Regardless, we believe that the 1% ITS medium could represent a useful tool for a more comprehensive follow-up investigation of the role of DNFA and DNCS in cancer proliferation and survival.

Our data indicate that insulin promotes both SREBP1 processing and transcription. Nuclear SREBP1 was elevated in cells cultured in 1% ITS and 0% FBS media, compared to 10% FBS medium. However, cytoplasmic SREBP1 precursor levels remained similar in both 1% ITS and 10% FBS medium conditions. This could indicate that SREBP1 precursor levels are maintained regardless of whether a portion of SREBP1 has been processed and migrated into the nucleus. Combining these observations with the gene expression data, we believe that insulin also has a stimulatory effect on *SREBF1* transcription in cancer cells, in agreement with previous findings from liver studies (22, 38).

DNFA is necessary for cell survival even in quiescent cancer cells that do not need membrane lipid synthesis for proliferation. In those cells, fatty acid oxidation (FAO) pathway hyperactivity has also been observed (69). DNFA and FAO are antagonistic lipid metabolism pathways that compose a futile cycle in cancer cells, but may represent a metabolic adaptation to promote cell survival under adverse (lipid-depleted) conditions (70). DNFA primarily relies on cytosolic NADPH derived from the pentose phosphate pathway (PPP) by glucose-6-phosphate dehydrogenase (G6PD), as well as malic enzyme (ME) and isocitrate dehydrogenase (IDH1), all of which are direct targets of SREBP1 (71). Cancer cells frequently promote NAPDH synthesis to support production of cellular anti-oxidants (e.g. glutathione) to counter elevated reactive oxygen species (ROS). These metabolic adaptations together may then represent a pro-survival mechanism, dependent on SREBP1.

## Conclusions

In summary, we have identified and validated a serum-free and insulin supplemented (ITS) medium condition that is well suited for controlled study of lipogenic gene activation and its mechanism of action in melanomas, and perhaps other cancer cell types.

## Author Contributions

Conceived and designed the experiments: SW and AMN. Performed the experiments: SW. Analyzed the data: SW and AMN. Wrote the paper: SW and AMN.

## Supporting information

**S1 Fig.**
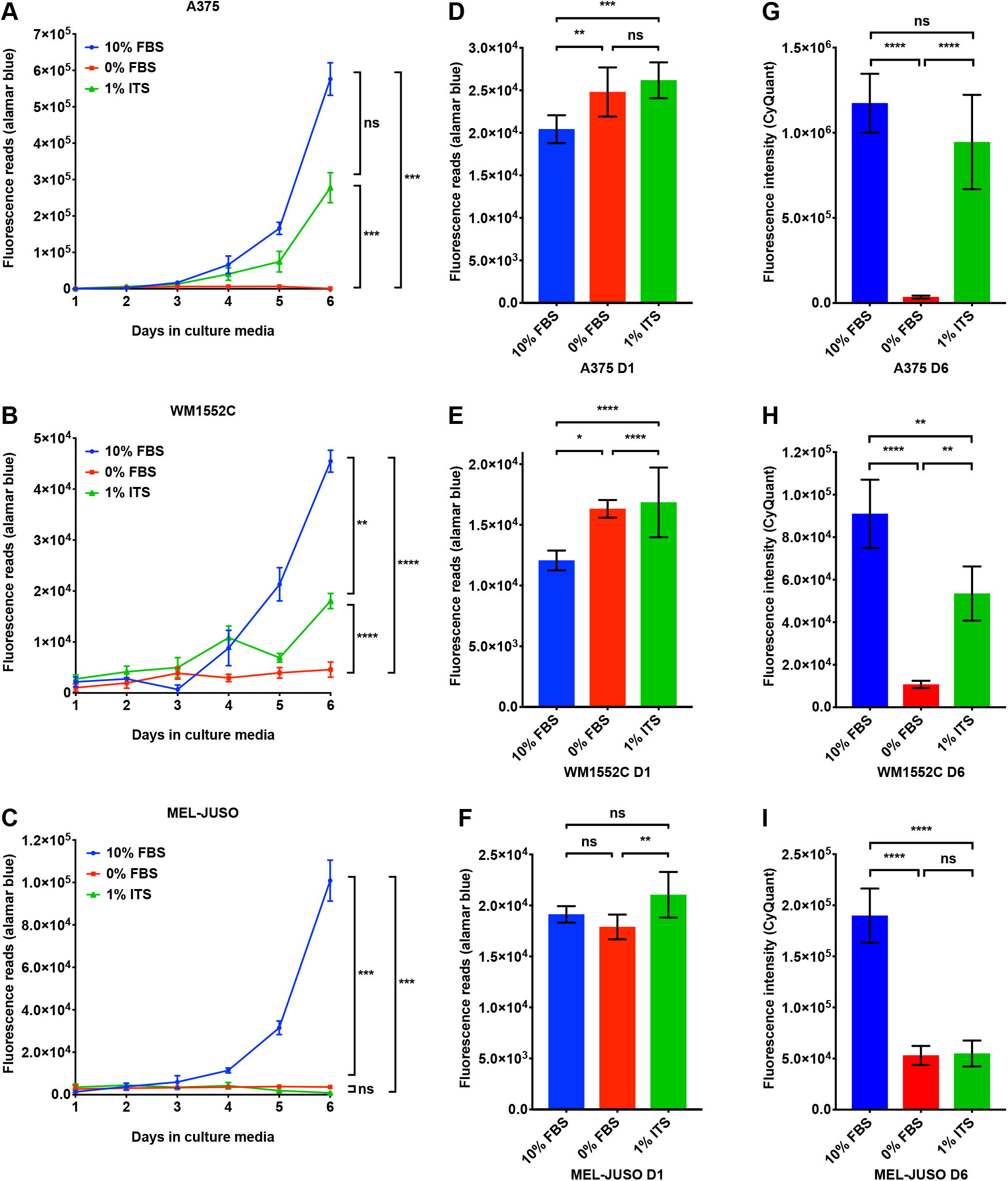
A lipid-free and insulin-supplemented medium supports proliferation of A375 and WM1552C, but not MEL-JUSO cell lines. (A – C) A375, WM1552C and MEL-JUSO cell lines were seeded in 10% FBS medium at day zero. On day one, cells were washed with PBS and transferred to the indicated medium conditions. Cell proliferation was measured with alamarBlue assay daily. Each data point represents the mean ±SD of quadruplicate samples. The results were analyzed using two-way repeated measures ANOVA followed by *post hoc* Tukey’s multiple comparison tests. (A) A375 cells: for culture time, F = 865.8, P < 0.0001; for culture condition, F = 282.8, P < 0.0001; for interaction between culture time and condition, F = 278.7, P < 0.0001. (B) WM1552C cells: for culture time, F = 351.1, P < 0.0001; for culture condition, F = 172.8, P < 0.0001; for interaction between culture time and condition, F = 137.4, P < 0.0001. (C) MEL-JUSO cells: for culture time, F = 501.6, P < 0.0001; for culture condition, F = 475.9, P < 0.0001; for interaction between culture time and condition, F = 584.5, P < 0.0001. (D – F) A375, WM1552C and MEL-JUSO cells were seeded in 10% FBS medium at day zero. On day one, cells were washed with PBS and changed to the indicated medium conditions. alamarBlue assay was performed on the cells cultured in the indicated medium for one hour. Results were analyzed using one-way ANOVA followed by *post hoc* Tukey’s multiple comparison tests. (D) A375 cells, F = 13.24, P = 0.0003. (E) WM1552C cells, F = 17.31, P < 0.0001. (F) MEL-JUSO cells, F = 6.985, P = 0.0061. Significant differences between medium conditions are indicated as *P < 0.05, **P < 0.01, ***P < 0.001 and ****P < 0.0001. ns, not significant. (G – I) A375, WM1552C and MEL-JUSO cell lines were seeded in 10% FBS medium at day zero. On day one, cells were cultured in the indicated medium conditions. CyQuant assays were performed on the cells cultured in the indicated medium at day six. Each data bar represents average measurement of five replicate samples. Results were analyzed using one-way ANOVA followed by *post hoc* Tukey’s multiple comparison tests. (G) A375 cells, F = 51.01, P < 0.0001. (H) WM1552C cells, F = 60.11, P < 0.0001. (I) MEL-JUSO cells, F = 96.82, P < 0.0001. Significant differences between medium conditions are indicated as *P < 0.05, **P < 0.01, ***P < 0.001 and ****P < 0.0001. ns, not significant.

**S2 Fig.**
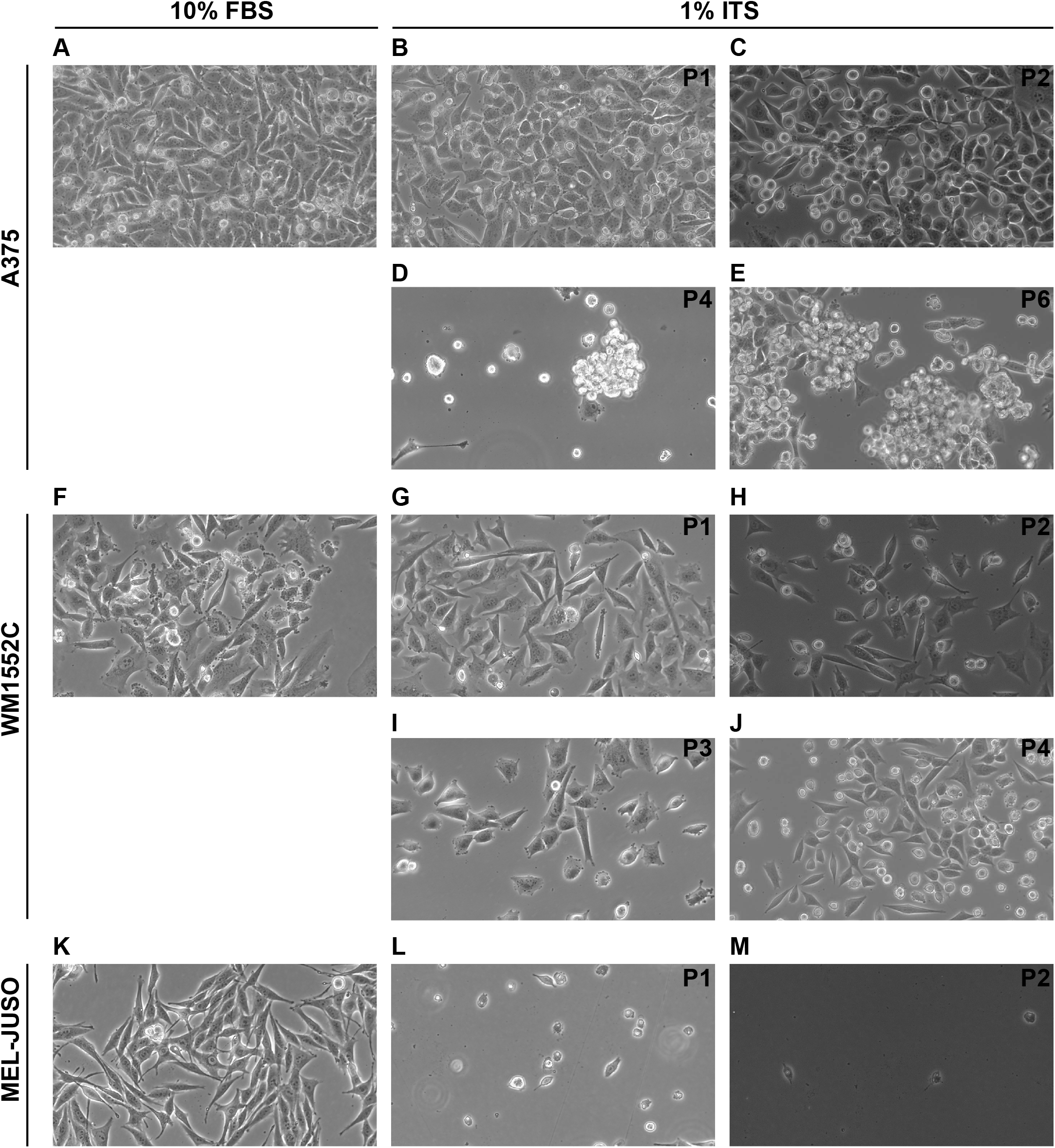
A lipid-free and insulin-supplemented medium supports long-term proliferation of melanoma cell lines A375, WM1552C but not MEL-JUSO. (A, F, K) A375, WM1552C and MEL-JUSO cell lines were routinely maintained in RPMI medium with 10% FBS. Morphologies of cells were recorded by light microscopy. (B-E) Morphologies of A375 cells cultured in 1% ITS medium from passage one (P1) to passage six (P6) over the course of six weeks were monitored by light microscopy. (G-J) Morphologies of WM1552C cells cultured in 1% ITS medium from passage one (P1) to passage four (P4) over the course of six weeks were monitored by light microscopy. (L-M) Morphologies of MEL-JUSO cells cultured in 1% ITS medium from passage one (P1) to passage two (P2). MEL-JUSO cells failed to proliferate in 1% ITS medium and could not be passaged. Morphologies of live cells were recorded by light microscopy with 40 × objective and 10 × ocular lens.

**S3 Fig.**
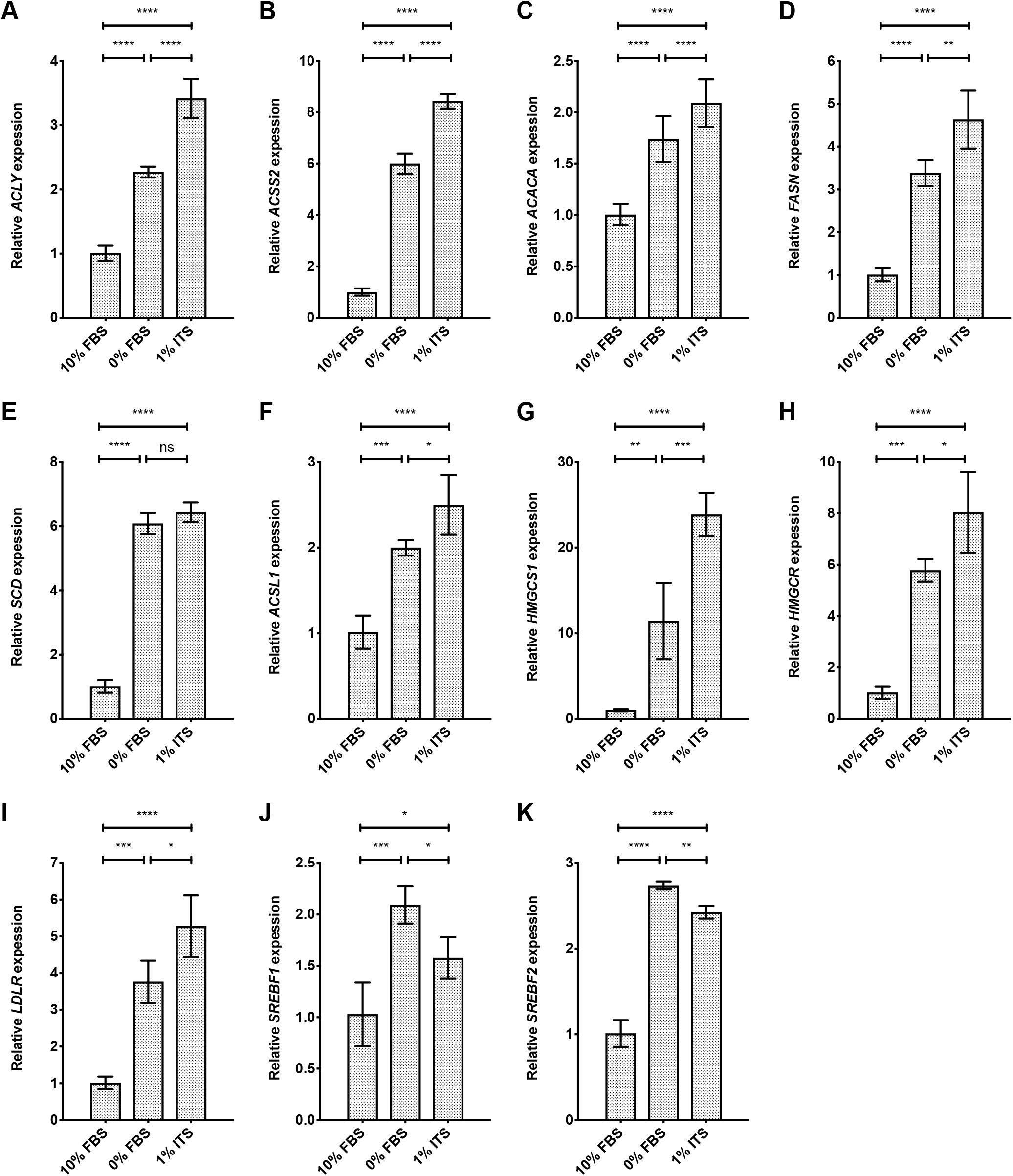
Lipid depletion combined with insulin supplement yields increased expression of lipogenic genes in MEL-JUSO cells. (A-K) The expression level of DNFA and DNCS genes was analyzed by RT-qPCR assay. MEL-JUSO cells were cultured in 10%, 0% FBS or 1% ITS medium for 24 hours. Significant differences between medium conditions are indicated as *P < 0.05, **P < 0.01, ***P < 0.001 and ****P < 0.0001 using one-way ANOVA followed by *post hoc* Tukey’s multiple comparison tests. ns, not significant. Each data point represents the mean ±SD of results from quadruplicate samples.

